# Genetic complementation reveals structure-function links in nodavirus RNA replication complex crowns

**DOI:** 10.64898/2025.12.10.693615

**Authors:** Johan A. den Boon, Helena Jaramillo-Mesa, Mark Horswill, Adam Jochem, Megan Bracken, Hong Zhan, Paul Ahlquist

## Abstract

Positive-strand RNA viruses replicate their RNA genomes in virus-induced, membrane-bounded organelles. As first found for nodaviruses, the necked cytosolic portals of these organelles bear ringed “crown” complexes of viral RNA replication proteins that drive synthesis, capping and release of new RNA genomes. Nodavirus crowns contain two 12-mer rings of viral protein A with C-proximal polymerase domains stacked. In the basal ring, protein A’s N-proximal RNA capping domains form a central, toroidal floor, while in the apical ring these domains extend radially outward. A third protein A conformation provides a putative central Pol domain interacting with the viral dsRNA replication intermediate in a vesicle beneath the crown. Protein A’s multiple conformations likely differentially contribute to crown assembly, RNA template recruitment, (−) and (+) strand synthesis, RNA capping, and progeny RNA release. Protein A’s high copy numbers may provide robustness to these processes. To test such concepts, we combined mutational, complementation and functional analyses. Strong complementation between null mutants in protein A’s polymerase and RNA capping active sites showed that they operate in independent protein A copies, likely at distinct sites. Thus, neither function is required in all protein A copies, nor are both required in any single copy. Lack of complementation between mutants in distinct RNA capping steps implied that major RNA capping steps must be performed in the same protein. Although RNA polymerase and capping activity were not required in all protein A subunits, none of a series of deletions across these domains were complementable, showing the importance of structural and other requirements for crown assembly, etc.. Surprisingly, RNA replication was more sensitive to depleting the fraction of subunits retaining protein A’s C-terminal intrinsically disordered region than polymerase or capping activity. These and other results reveal and illuminate the cooperative, interdependent nature of protein A’s diverse functions.

**Author summary:** Positive-strand RNA viruses represent the largest genetic class of viruses and include human, animal, and plant pathogens causing major agricultural, economic, and environmental consequences. Using no DNA intermediates to multiply their RNA genomes, these viruses modify cellular membranes into novel, infection-specific RNA replication organelles. Emerging results show that RNA replication proteins encoded by many or most of these viruses assemble into ringed, crown-like viral protein complexes gating portals to these compartments. We previously revealed that nodavirus crowns contain two stacked 12-mer rings of viral replicase protein A, which contains polymerase, RNA capping and other domains. The nodavirus experiments reported here are among the earliest explorations in cells to illuminate the functions and interactions of such multi-domain RNA replication proteins in the context of their highly multimeric crowns. Critical questions include whether all domains and interactions are required in all protein A conformations, whether protein A multiplicity might provide dose-responsive redundancy for any crown functions, or whether defects in individual protein copies might inhibit or even poison operation of the entire crown. The results have significant implications for positive-strand RNA virus biology and thus for efforts toward virus control and beneficial uses.

## INTRODUCTION

Positive-strand RNA ((+)RNA) viruses are an extensive, diverse class of pathogens that pose major risks to public health. They include coronaviruses like pandemic SARS-CoV-2, alphaviruses like chikungunya virus (CHIKV), flaviviruses like dengue virus and tumor virus hepatitis C virus, picornaviruses like poliovirus, “cruise ship” noroviruses, and many others. The core of their life cycle and the focus of most of their genetic coding capacity is genome replication, which cycles between mRNA-sense single stranded (ss) RNA genomes that are substrates for translation, replication and encapsidation, and double stranded (ds) RNA intermediates sequestered in virus-induced vesicles [1–4].

Nodaviruses are a family of (+)RNA viruses that infect vertebrate and invertebrate hosts [5,6]. The best-studied member, flock house virus (FHV), has become an advanced model for (+)RNA virus genome replication and other processes. FHV’s 4.5 kb genome is divided between 3.1 kb RNA1 and 1.4 kb RNA2. RNA1 is translated to produce protein A, a 998 aa multifunctional RNA replication protein (Figure 1A). Protein A’s C-proximal half contains an RNA-dependent RNA polymerase domain, Pol, linked via a 57-aa highly flexible linker to an N-terminal half containing two membrane interaction domains and a 5’ RNA capping domain. RNA1 also encodes 0.4 kb subgenomic RNA3 that expresses RNAi suppressor B2 and, in an alternate frame, B1. B1 coincides with the C-terminus of protein A, a poorly conserved region whose function is not well understood. RNA2 encodes the viral capsid protein.

**Figure 1.**
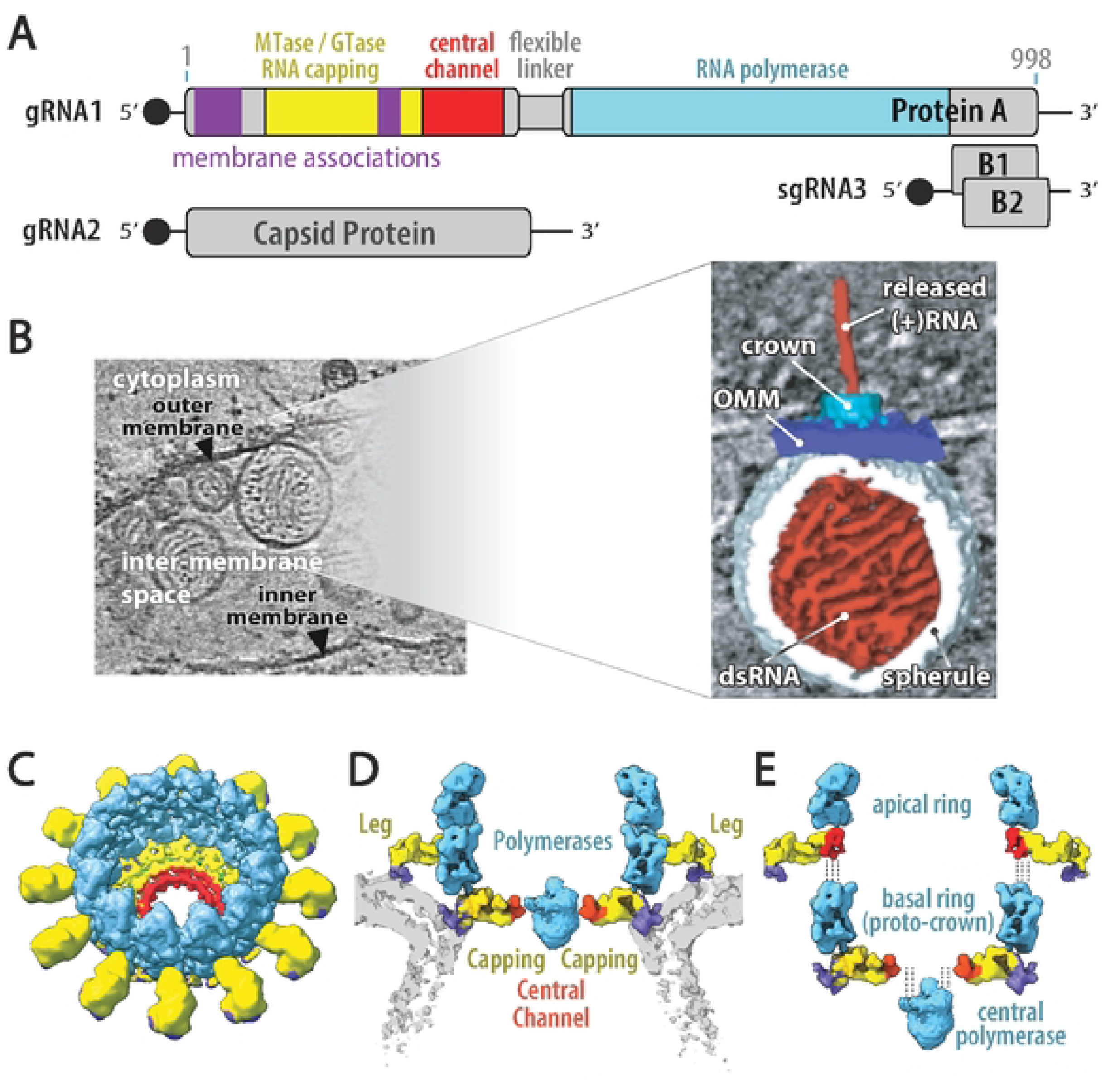
Organization of nodavirus RNA genome and crown RNA replication complexes. (A) The bipartite nodavirus RNA genome encodes 112 kDa / 998 aa protein A, the multifunctional RNA replication protein, on genomic RNA1 and capsid protein on genomic RNA2. Protein A’s (-terminal half contains an RNA-dependent RNA polymerase domain (cyan and its N-terminal half containsa S’RNAcapping domain (yellow),a“central channel”domain (red), and membrane interaction domains (violet). A 57 aa long flexible linker (gray) separates the polymerase and central channel domains. During RNA1 replication, a subgenomic RNA3 is produced that encodes two small proteins: protein B1 of unknown function (co-terminal to protein A) and protein B2 that counteracts antiviral RNAi defense mechanisms. (B) Upon infec-tion, to organize and protect the RNA replication process, protein A targets itself to the outer mitochondrial membrane, recruits viral RNA templates, induces spherular membrane invaginations to compartmentalize the dsRNA replication intermediates, and multimerizes to form a 12-fold symmetric crown structure that gatesthe spherules’ necked aperture to the cytosol.(Cl Perspective, (D) cross-section and (E) partially disassembled viewsof the nodavirusmultimeric protein A crown structure. As shown, the crown contains two stacked 12-mer ringsof viral protein A, whose domains are color coded as in the genome map of panel A. 12-mer apical and basal rings of polymerase domains (cyan) stack to form a central turret, with the associated N-terminal protein A regions(yellow/red/violet) in two different conformations.In the basal ring, which constitutes a “proto-crown” early assembly intermediate, the N-terminal regions assemble into the crown’s floor, whose central opening accommodates an additional copy of the RNA polymerase adjacent to the dsRNA replication intermediate inside the spherule vesicle beneath (not shown in the panel Cperspective, to provide an unob-scured view of the floor and central channel). In the apical ring, the N-terminal domains project radially outward as membrane-interacting legs.In panel D, the lipid bilayer of the replication vesicle neck is shown in light gray.

Using FHV and cryo-electron microscopy (cryo-EM), we showed that (+)RNA virus genome replication proteins are organized in 12-fold symmetric, double-ringed “crowns” that gate the neck of invaginated RNA replication vesicles (“spherules”) containing the dsRNA replication intermediates (Figure 1B) [7,8]. Subsequent findings showed that alphaviruses form similar 12-fold symmetric crowns on their RNA replication vesicles and that coronaviruses also make membrane-spanning, RNA replication-linked crowns sharing many underlying principles [1,2,9–13].

FHV’s RNA replication occurs on outer mitochondrial membranes (OMM) (Figure 1B), where protein A first forms a membrane-linked proto-crown – a 12-mer protein A ring that becomes the basal ring of the mature crown (Figure 1CDE, [14]). Immediately C-proximal to the N-terminal RNA capping domain, sequences previously known as the “iceberg region” [15] assemble a central channel docking site for a central Pol domain actively synthesizing (+)RNAs. Subsequent closely-linked steps include recruiting (+)ssRNA viral genomic RNA templates, adding a second 12-mer protein A apical ring atop the proto-crown, synthesizing complementary viral (-)RNA and depositing the resulting dsRNA replication intermediate in a spherular OMM invagination immediately below the crown [1,3,14,16–18]. The resulting mature RNA replication complex repeatedly copies the dsRNA replication intermediate to synthesize and 5’ cap new (+)ssRNA genomes that are released to the cytosol to be translated, form new replication complexes or be encapsidated into virions.

Mature, double ring nodavirus crowns on active RNA replication complexes are composed of protein A in three conformations [14] (Figure 1CDE). The basal ring, whose structure is indistinguishable from the proto-crown precursor, consists of a circle of 12 Pol domains with their N-proximal membrane-binding / RNA capping / central channel domains interacting to form a highly ordered toroidal floor below. In the apical ring a similar circle of 12 Pol domains is stacked atop the first, with their N-proximal domains again below but separated and extended outward. The resulting apical and basal rings of outer membrane interacting domains likely tightly constrain the strong curvature of the spherule membrane neck.

The central portion of the floor in mature, double ring crowns is occupied by a density absent in proto-crowns (Figure 1D). While no atomic or near atomic structure has been determined for this central density, multiple points strongly implicate it as the active polymerase for (+)RNA synthesis, including its optimal positioning to read the vesicle-sequestered viral dsRNA and release nascent RNA products, juxtaposition to the floor RNA capping domains and comparable size and shape to the proto-crown pol domain [8,14]; equivalent positioning of the sole nsP4 RNA polymerase in CHIKV crowns [19]; and inaccessibility of the 24 pol domains in the crown central turret to the dsRNA template sequestered below the crown floor [1,8]. Such a Pol domain might have detached from the apical or basal ring and use the long flexible linker to protein A’s N-terminal half to reach the central channel. Alternately, the central density might be a 25^th^ Pol domain, either a rare copy expressed or processed free of linkage to protein A’s N-proximal domain, or part of a full-length protein A whose unique, flexible membrane anchorage has not yet been visualized.

Protein A’s multiple roles, copies and conformations raise critical questions about how different conformers may be specialized for particular functions and how these conformers cooperate for crown assembly and operation. Under different scenarios posed above, e.g., a central Pol domain might be part of a specialized, full-length protein A that supplies and integrates all Pol and RNA capping activities, or might provide only Pol activity while depending for capping on another protein A conformation, such as the basal ring providing the adjacent crown floor. Further questions include whether all domains, interactions, trafficking signals and membrane binding motifs are required in all protein A conformations, or if in certain contexts some might be dispensable or supplied *in trans* by other copies of protein A. Moreover, it is presently unknown to what extent the multiplicity of protein A copies in the crown provides dose-responsive redundancy and robustness for any functions, or whether defects in individual proteins might interfere with adjacent interaction partners or even poison operation of the entire crown.

To address such questions, we combined genetic complementation with other approaches to separate specific functions between distinct, co-expressed copies of protein A, and assayed the effects on RNA replication and selected sub-steps. The results identify *trans*-separable and *cis*-inseparable functions of protein A and provide further insights into the roles, independence and interaction of multiple protein A states in crown operation and viral replication in general.

## RESULTS

### DNA-launched RNA replication

To expedite replication studies of wt and engineered derivatives of nodavirus RNAs, we launched RNA replication from transfected DNA expression plasmids in S2 cells of *Drosophila melanogaster*, a permissive host for natural infection (Figure 2A). Since the nodavirus capsid protein is not required for RNA replication, our standard transfection assay omitted RNA2 and was based on replication of plasmid-launched RNA1. Since RNA1 encodes the sole nodavirus RNA replication protein, protein A (Figure 1A), plasmid launching of wt RNA1 *cis-*replication would complicate determining whether introduced mutations affected protein A function or RNA1 template function or both. To overcome this, we used a “*trans*-replication” system in which protein A and a modified RNA1 replication template were expressed from separate plasmids. To block its function as a replication template, the protein A mRNA lacked 5’ and 3’ *cis*-replication elements of RNA 1 and was also recoded at translationally silent positions across an RNA region previously found important for template recruitment [17]. In parallel, the RNA1 template (RNA1fs) was rendered deficient for protein A translation by a 5’ proximal frameshift mutation and, as an additional precaution, a D692-to-E mutation of the polymerase catalytic site. As shown in Figure 2B, co-expressing wt protein A and RNA1fs (lane 3) replicated RNA1fs to levels far above basal background of plasmid-transcribed protein A mRNA (lane 1) or RNA1fs (lane 2), and produced even higher levels of sgRNA3, which lacked any DNA-derived transcript background and thus provided a true measure of RNA replication and transcription.

**Figure 2.**
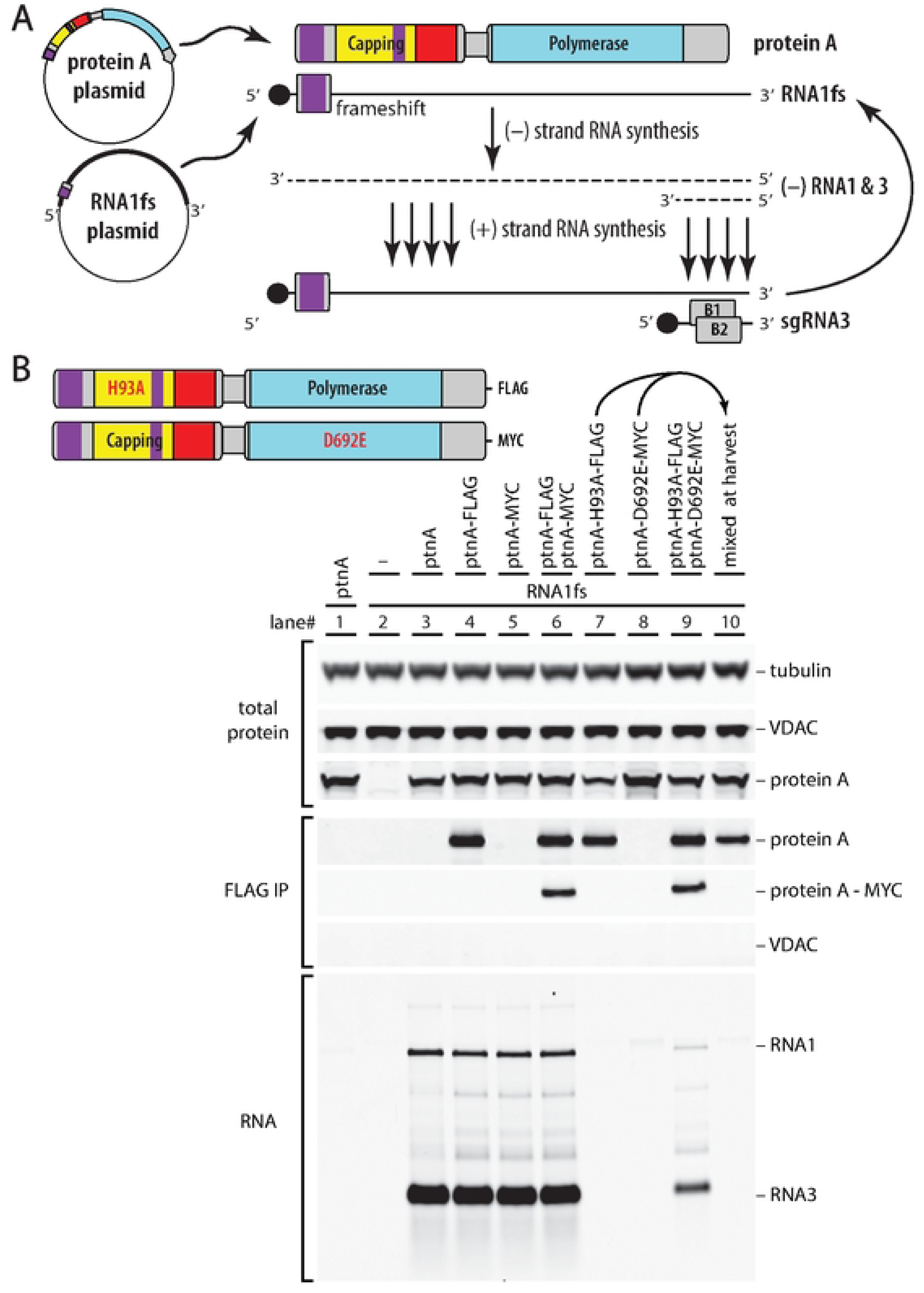
Nodavirus DNA-launched trans-replication system reveals complementation between RNA capping and RNA polymerase mutants. (A) A dual DNA plasmid system allows separate genetic manipulation of protein A functions and RNA template functions.Oneplasmid launches expression of protein A from anon-replicable mRNA and a second plasmid transcribes areplicable RNA1 template rendered incapable of translating protein A via an early frameshift mutation (RNAlfs).The resulting protein A uses the RNAlfs transcript in trans as a template for RNA replication and transcription, through a single round of(-) strand synthesis and subsequent repeated rounds of(+) strand synthesis. During these processes, production of a partial (-) strand launchesreplication of the small subgenomic RNA3.Being solely derived from RNA replication and never transcribed from DNA. RNA3 provides a direct measure of RNA replication activity, without needing to correct for any DNA-derived background. (B) Plasmid-launched assays of the RNA replication and complementation activities of wt protein A and active site null mutants in its RNA capping and polymerase domains, and immunoprecipitation assays of interaction between (-terminal FLAG- and MYC-tagged protein A variants as illustrated at the top left. Western blots within the uppermost bracket show total levels of nodavirus RNA replication protein A, tubulin as a loading control and VDAC as a control for mitochondrial outer membrane proteins. In the intermediate bracket, western blots show the levels of total protein A,MYC-tagged protein A, and VDAC after FLAG immunoprecipitation. At the bottom,a northern blot shows accumulation of nodavirus RNA1 and RNA3 replication products after co-transfection of plasmids expressing the RNA1fs replication template and the wt, taggedand/or mutant variant(s) of protein A indicated above each lane. All experiments were performed three or more timesand representative blots are shown.

### RNA capping and polymerase protein A mutants jointly multimerize in vivo to complement each other’s defects

As outlined in the Introduction, protein A’s multiple functions in RNA synthesis and capping might be executed by a single preferential protein A copy or dispersed among interdependent action of some subset of the 24-25 protein A copies in mature RNA replication crowns. Accordingly, in a first genetic complementation experiment, we co-expressed an RNA capping-null protein A with catalytic site mutation H93A with an RNA polymerase-null protein A with catalytic site mutation D692E to assess their capacity to rescue each other’s defects for replicating the RNA1fs template. H93A alters the proposed site of m^7^GMP covalent esterification to protein A for subsequent 5’ capping of RNA1 and RNA3 [15,20,21] while D692E alters the second position in the highly conserved GDD motif essential for RNA-dependent RNA polymerase function [22–24]. Notably, while the individual mutants failed to support RNA replication as expected (Figure 2B, lanes 7-8), their co-expression restored RNA3 accumulation (lane 9). For simplicity, hereafter such mutant protein A derivatives are referred to by their genotype, e.g., H93A or D692E.

Thus, protein A’s RNA polymerase and RNA capping functions can be executed by separate protein copies. Such complementation would appear to require interaction of the distinct mutant proteins in a single crown, since only one such crown sits above each RNA replication vesicle and its enclosed viral dsRNA, copies this dsRNA template and releases the resulting nascent (+)RNA products to the cytosol (Figure 1B). To test for such interaction by co-immunoprecipitation (co-IP), this experiment used protein A derivatives carrying small C-terminal epitope tags. As controls, FLAG- and MYC-epitope tagged versions of wildtype protein A showed that these small C-terminal tags do not significantly interfere with RNA replication (Figure 2B, lanes 4-6). Anti-FLAG antibody-mediated co-IP followed by anti-MYC antibody western blot detection showed that, when co-expressed in vivo, differentially tagged wt protein A’s (lane 6) or the two tagged mutants (lane 9) physically interact, leading to pull down of the MYC-tagged protein, which did not occur when the wt or mutant MYC-tagged protein was expressed alone (lanes 5 and 8). Such interaction and co-IP only occurred when cells co-expressed both proteins in vivo (lanes 6 and 9) and not when cell extracts from single protein-expressing cultures were mixed post-harvest (lane 10). Further, co-IP of H93A and D692E mutant proteins was not mediated indirectly by individual association of proteins with hypothetical remnants of outer mitochondrial membrane that somehow survived the detergent conditions used, since another prominent outer mitochondrial membrane protein, VDAC, was not pulled down under any conditions tested (Figure 2B). Accordingly, the complementing mutant proteins interact in stable protein complexes, consistent with the expectation that they function jointly in individual crowns.

### Sequential RNA capping steps are carried out within the same protein A subunit

Despite extensive sequence divergence, nodavirus protein A and alphavirus nsP1 RNA capping domains share a closely superimposable methyltransferase fold [14], assembly into 12-mer rings [8,10,11,14,19], and an unusual capping pathway. While cellular and many viral RNA capping pathways add GMP to a 5’ ppNpN… RNA end and then methylate GMP at N7, protein A and nsP1 first N7 methylate GTP, form a covalent link with the resulting m^7^GMP, and transfer that m^7^GMP to 5’ diphosphorylated viral RNA [20,21,25–27]. Critical residues in the protein A RNA capping domain include aspartic acid D141, which corresponds to CHIKV nsP1 D37 that coordinates methyl donor S-adenosyl-methionine for the early N’7 GTP methylation step, and the above-mentioned histidine H93, which corresponds to CHIKV nsP1 H89, to which m^7^GMP is later esterified [15].

To further test the relation of these RNA capping steps to each other and to viral RNA polymerase action, complementation tests were performed among untagged protein A capping mutants H93A and D141A and polymerase active site mutant D692E (Figure 3). Varied ratios of the mutant protein A expression plasmids were included to explore possible effects on functional complex assembly and complementation. As shown in Figure 2B, lane 7 and Figure 3A and 3C, lane 4, the H93A protein A mutation blocked detectable accumulation of RNA replication products, as did D141A (Figure 3B, lane 4 and 3C, lane 14) and D692E (Figure 2B lane 8 and Figure 3A and 3B, lane 14). Extending the Figure 2B results, H93A and D692E complemented each other detectably across all plasmid ratios tested, from 9:1 to 1:9, with a broad optimum slightly skewed toward more plasmid DNA for the H93A capping mutant (Figure 3A lanes 7-9). Similarly, the D141A RNA capping and D692E polymerase mutants complemented each other across all tested ratios, with a broad optimum near balanced DNA levels (Figure 3B, lanes 8-10).

**Figure 3.**
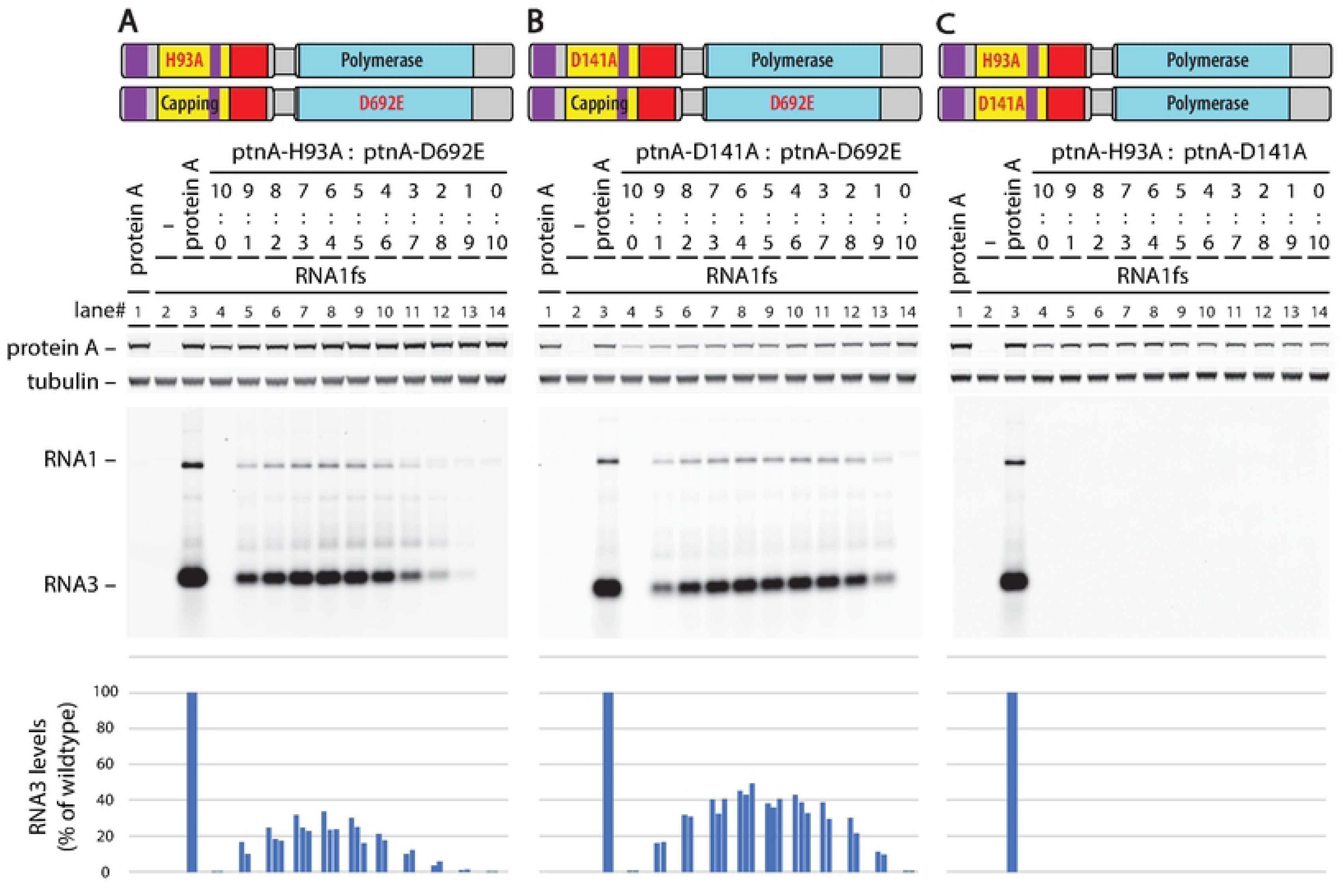
Protein A nullmutants in sequential RNA capping steps each trans-complement an RNA polymerase nullmutant but not each other. (A) and (BJ Plasmid-launched transreplication assays as in Fig. 2A show that co-expressing either of two protein A mutants catalytically inactive in distinct RNA capping steps ((A) H93A or (Bl 0141A) with a catalytically inactive RNA polymerase mutant (D692E) rescues RNA replication of the RNA1 fs template and its subgenomic RNA3. The indicated wide range of plasmid ratios(9:1, 8:2,etc.) wasused to test for possible differential sensitivity of replication to depletion of one or the other complementing protein A allele. Representative blotsare shown and the results of three independent experiments are plotted in the bar graphs at the bottom. (Cl In parallel experiments, when co-expressed, the two RNA capping null mutants (H93A and 0141A) are unable to trans-complement each other to restore RNA replication. Again,a representative blot isshown and the results of three independent experiments are plotted in the bar graph at the bottom.

Notably, capping mutants H93A and D141A did not complement each other at any tested ratio (Figure 3C). This inability to be *trans*-complemented implies that the sequential RNA capping steps of GTP methylation (disrupted in the D141A mutant) and m^7^GMP transfer to the 5’ end of the RNA (disrupted in the H93A mutant) need to be carried out by a single protein A molecule, and that the m^7^GTP intermediate, e.g., does not diffuse between two neighboring protein A molecules in the crown. This functionally confirms expectations from the relatively close positioning of the residues involved in nodavirus protein A and Chikungunya nsP1 and inferences that dynamic movements within a single subunit mediate m^7^GTP transfer to the m^7^GMP esterification site [14,28].

### Protein A’s C-terminal intrinsically disordered region is required at high copy numbers

The Figure 2 and 3 results show that protein A’s Pol domain functions when linked to an inactive capping domain and its capping domain functions when linked to an inactive Pol. Accordingly, it seemed possible that the complementing ability of the intact Pol or capping domain might also be retained after deleting some sequences from the alternate, inactivated domain. As an initial test, seventeen 100 aa deletions tiled at 50 aa intervals (Figure 4A) were assayed for their ability to complement capping- or polymerase-defective mutants H93A and D692E. To facilitate mitochondrial delivery and association, all deletions retained the N-proximal transmembrane domain (aa 9-47) that targets protein A to the outer mitochondrial membrane [29]. When assayed for complementation at equal plasmid levels (Figure 4B), none of these deletions detectably complemented the D692E Pol mutant, and the H93A capping mutant was only complemented at extremely weak levels (∼1% of wt protein A) by a single mutant, C-terminal deletion Δ897-988. Thus, both the Pol and RNA capping activities require a largely complete protein A context for effective expression.

**Figure 4.**
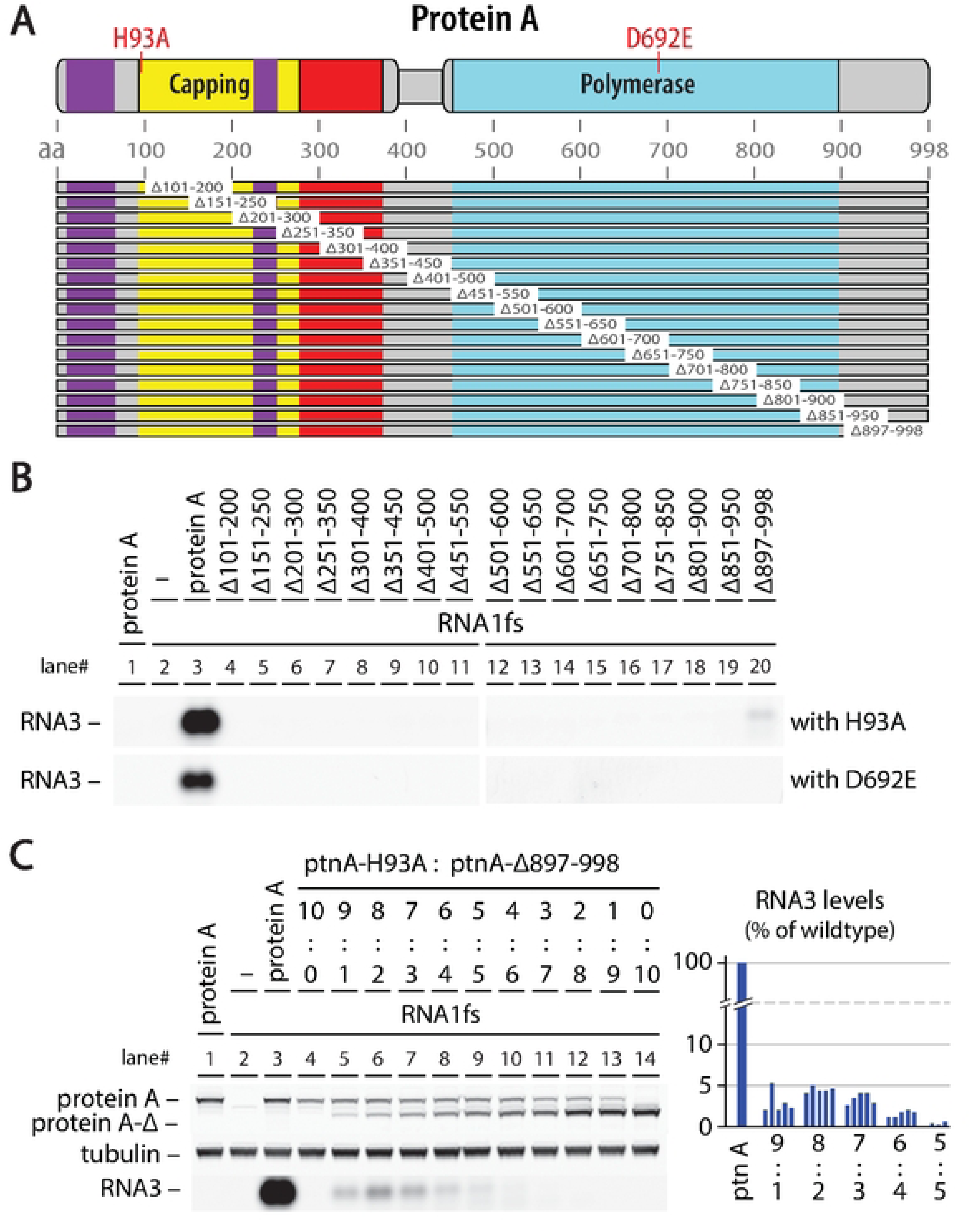
Although RNA capping and Pol functions arenot required in all protein A subunits, deletionsperturbing the struc-ture of either domain are not complementable. (A) Map of seventeen 100 aa deletions tiled across protein A. All deletions retained the first 100 aa to ensure their mitochondrial targeting and membrane insertion (Miller and Ahlquist 2002). (Bl Plasmid-launched trans-replication assays co-expressing wt protein A or the indicated deletion mutants from panel A with capping null mutant H93A (upper row) or polymerase null mutant D692E (lower row). All experiments were repeated at least three times and representative northern blots are shown. As illustrated, all deletions lost the ability to detectably complement mutant H93A or D692E, except for the final deletion .!1897-998, which very weakly (−1% of wt) complemented H93A but not D692E. (Cl Detailed analysis of complementation between capping active site null mutant H93A and .!1897-998, which deletes the poorly conserved, intrinsically disordered (-terminal regionof protein A. As in Fig. 2, a wide range of plasmid ratios(9:1, 8:2, etc.) was used to define the relative sensitivity of replication to depleting either complementing protein A allele. A representa-tive blot is shown and the results of five independent experiments are plotted in the bar graph at the bottom. As shown, in contrast to the relatively balanced complementation curves in Fig.3AB, in this case the maximum complementationefficiency of −5% wt RNA replication was achieved at a ratio of 8:2 between the plasmidsexpressing H93A and .!1897-998, respectively.

The region deleted in the Δ897-998 mutant that barely complemented H93A is predicted to be intrinsically disordered and its function is weakly restorable by substitution with certain nonviral sequences [30]. Intriguingly, when tested across a wider range of complementing plasmid ratios (Figure 4C), a higher maximum complementation of ∼5% of wt protein A activity was achieved at an optimal H93A:Δ897-998 plasmid ratio of 8:2. Thus, even though for this pair the Δ897-998 mutant was the only source of RNA capping activity, the best overall performance was achieved when the truncated protein was present at notably lower levels than the capping null H93A mutant which retains a full-length, wildtype C-terminal domain (see Figure 4C western blot). These results show that the C-terminal intrinsically disordered domain provides critical RNA replication function(s) that are more sensitive than protein A’s enzymatic functions to reduced copy numbers.

### Both of protein A’s membrane association domains are required in all protein A subunits

Protein A interacts with membranes through segments M1 and M2 ([14] and Fig 1A, D and E, purple domains). M1 spans the N-terminal aa 1-64, including the transmembrane targeting element that directs protein A or other proteins like GFP to the outer mitochondrial membrane [29]. M2 is a mostly alpha helical surface loop protruding from the RNA capping domain [14]. The protein A subunits in the apical and basal rings of the mature crown interact with membrane segments of very different curvature (Figure 1DE), while the flanking sequences and possible membrane connection of the central density putative active Pol domain (Figure 1DE and Introduction) are unknown. Moreover, protein A multimerization might allow one protein A derivative to support mitochondrial targeting of another. Accordingly, it is not known whether M1 and M2 are both required in any or all subunits of the apical or basal protein A rings or whether either is required in connection with the central density.

To address these questions, we made two mutants (Figure 5A): ΔM1 is a deletion of aa 9-64, while in ΔM2 the membrane-associating alpha helical loop (aa 236-251) was replaced by Gly-Gly-Pro-Gly-Gly to bridge the flanking portions of the RNA capping domain. Each mutation blocked RNA replication (Figure 5B, lanes 4 and 14). Accordingly, using the same broad ratio complementation design of Figure 3, ΔM1 and ΔM2 were tested for their capacity to complement each other or the H93A capping and D692E polymerase mutants. Figure 5B shows that none of these pairings supported RNA replication at any tested ratio, demonstrating that both membrane association determinants are required *in cis* for the RNA capping and Pol enzymatic functions and that they are needed on all protein A subunits in the crown.

**Figure 5.**
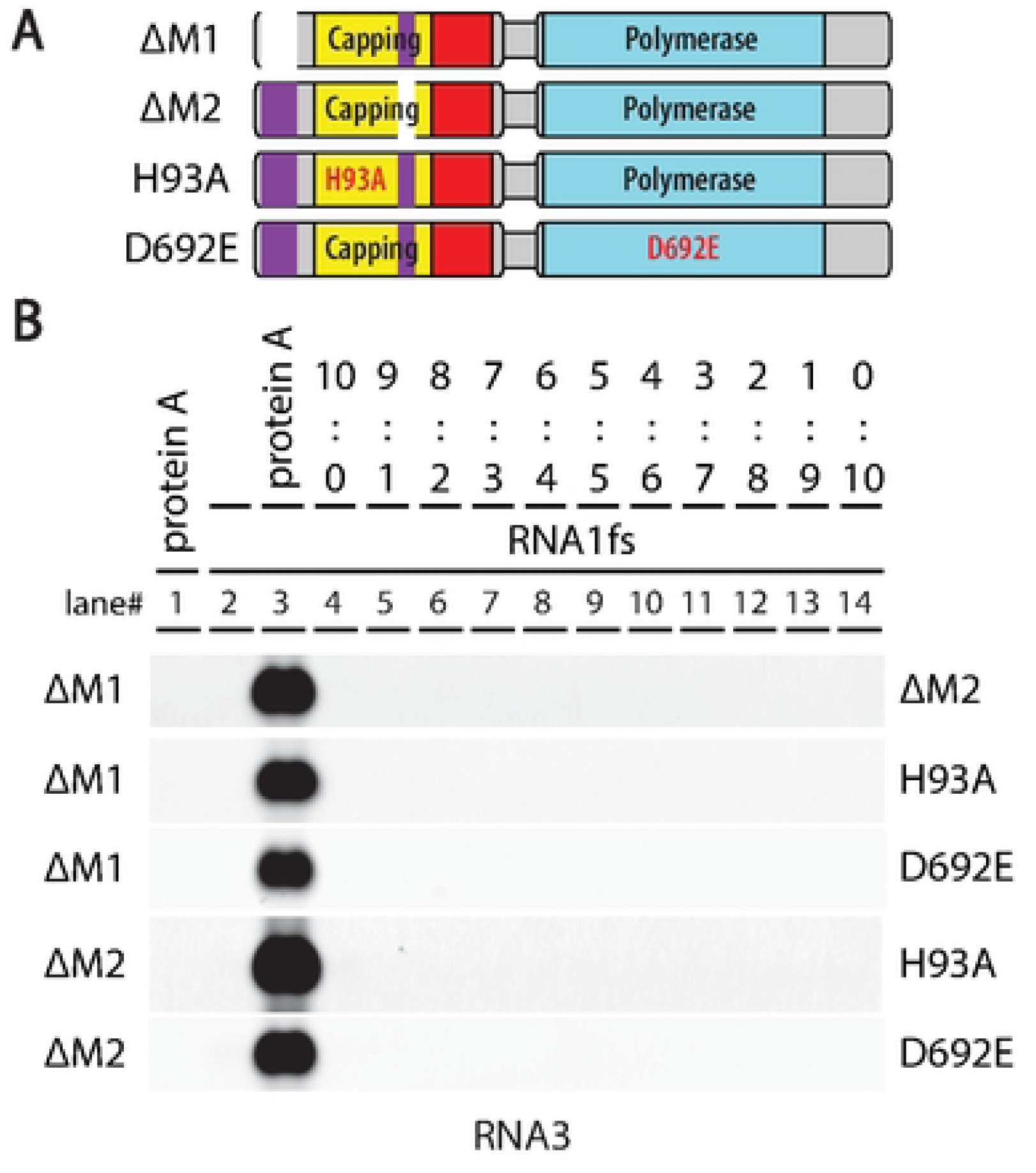
Both of protein A’smembrane association domains are required in all protein A subunits. (A) Map of targeted deletion of the membrane interaction domains in protein A and of the catalytically inactive capping mutant H93A and polymerase mutant D692E. The individual membrane domain deletion mutants did not complement each other, nor could either one of themcomplement capping null mutant H93A or polymerase null mutant D692E. Independent repeat experiments confirmed the results.

### Mutations at Pol-Pol interfaces inhibit RNA synthesis

The proto-crown floor, composed of protein A’s membrane-interaction, RNA capping and central channel domains (Figure 1D), is the most stable and well-ordered element in the nodavirus crown, and serves as a highly rigid foundation for proto-crown and full crown assembly [14]. By contrast, the ring of Pol domains in the proto-crown above this floor (the basal Pol ring in the full, double ring crown, Figure 1D) is much more flexible, with individual subunits rocking forward and backward on their basal contacts with the floor by ∼5° [14]. Consequently, whether these flexible lateral interactions between Pol subunits were essential for proto-crown assembly and other replication functions was unclear. To examine this, we selected for mutation three amino acid (aa) positions in loops on the Pol surface at the interface between proto-crown Pol domains: A677, R758, and T761 (Figure 6A). Mutants were generated using PCR with primers having degenerate sequences at these three codon positions and many though not all possible amino acid substitutions were obtained. Figure 6B and C show that RNA replication tolerated many mutations at aa 677 and 761, but that mutations A677K, T761E and T761P abolished RNA replication.

**Figure 6.**
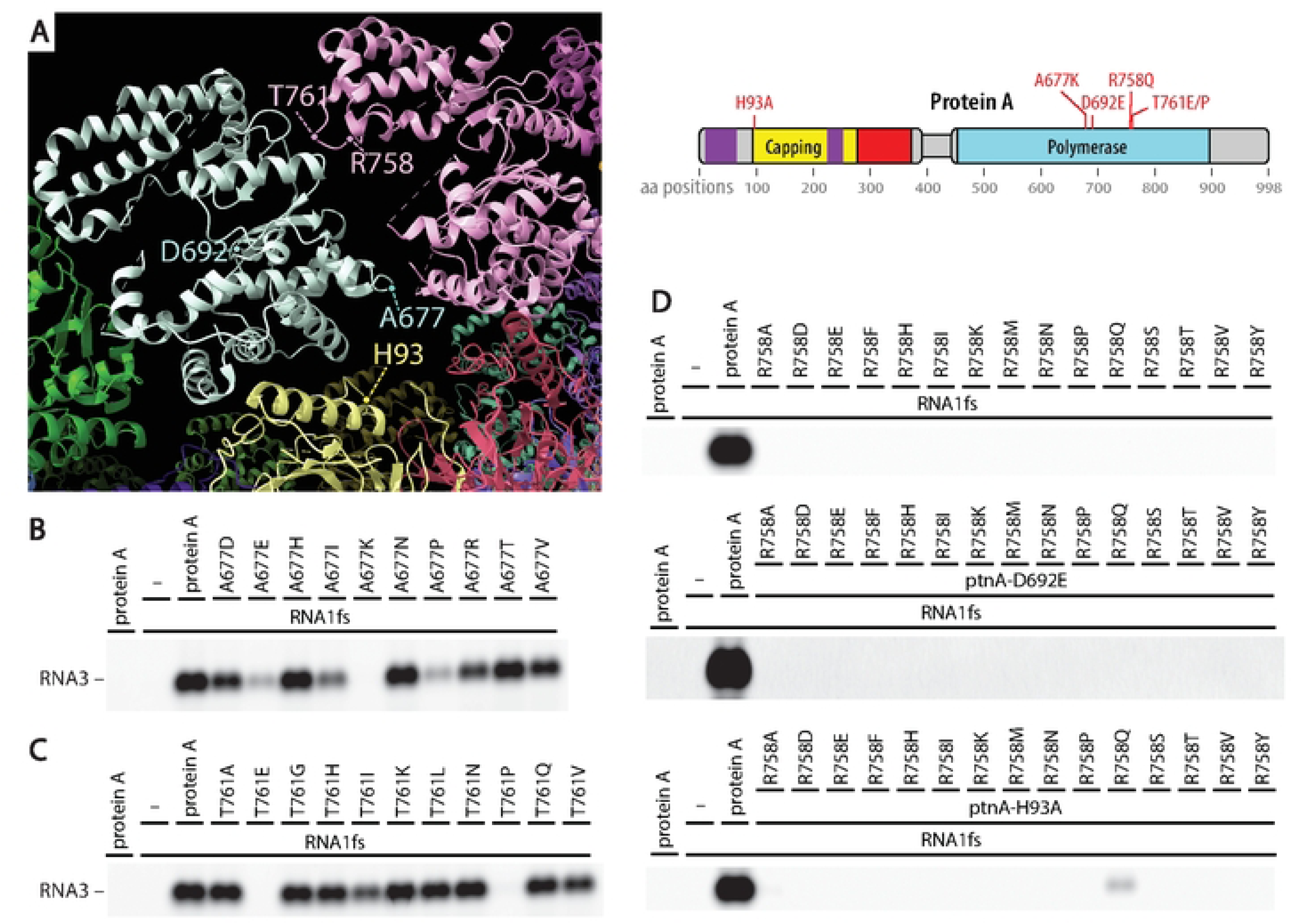
POL-POL interface positions tolerate varying degrees of mutation. (A) In the proto-crown structure, RNA polymeraseamino acids A677, R758 and T761 are positioned closest to the interface between polymerase subunits and can be expected to mediate lateral POL-to·POL interactions. Positions are also shown for the catalytic sites at H93 and D692 for RNA capping and RNA polymerase, respectively. Northern blots in B-D show levels of subgenomic RNA3 synthesis supported by panelsof mutants at these positions. Positions A677 (BJ andT761 (C) were mutation-tolerantexcept for changes from alanine to lysine and from threonine to either glutamate or proline, respectively.(D) By contrast, the R758 position did not tolerate any mutation other than from arginine to glutamine which retained some capacity to provide RNA capping activity. Mutants A677K, R758Q, T761E and T761P were chosen for follow-up experiments.

In contrast, at aa R758, all of 15 tested mutations blocked RNA replication (Figure 6D). To further examine the nature of these defects, each R758 mutation was tested for possible complementation with the protein A D692E Pol null and H93A RNA capping null mutants (Figure 6E). All tested mutations at aa R758 blocked complementation with the D692E Pol null (upper panel). Additionally, only mutations R758Q and R758A showed weak to faint complementation with mutation H93A in the distal RNA capping domain (lower panel). Thus, nearly all mutations in aa R758 blocked RNA polymerase and RNA capping activities, suggesting defect(s) in a fundamental function or functions underlying both enzymatic activities.

### Mutations at Pol-Pol interfaces inhibit proto-crown assembly

We previously showed that proto-crowns can be purified from cells and imaged by cryo-EM tomography [14]. Since cryo-EM tomography is too time-intensive to assay growing numbers of protein A mutants, we developed a more facile proto-crown imaging assay based on negative staining. To express protein A to consistent levels in all cells of a test population, recombinant baculoviruses were generated to express wt and desired mutant forms of protein A bearing C-terminal alfa tags that do not interfere with nodavirus replication and allow purifying intact proto-crowns from cells [14]. These baculoviruses - collectively expressing alfa-tagged wt protein A and its mutants H93A, D692E, A677K, R758Q, T761E and T761P (see Figure 6A-E) - were used to infect *S. frugiperdia* Sf9 cells and, 72 hr post infection, mitochondria were extracted and processed for proto-crown isolation as described [14]. The resulting extracts were spread on EM grids, negatively stained with uranyl acetate and imaged by transmission EM.

For extracts from cells expressing wt protein A, this negative stain assay revealed numerous ∼18-19 nm diameter rings (Figure 7A) resembling equivalent proto-crown preparations after flash freezing and imaging by single particle cryo-EM for structural analysis [14]. Further, just as for single particle cryo-EM analysis of proto-crowns, 2D image classification showed that these negatively stained structures included 12-mer and 11-mer rings (Figure 7A). Numerous cryo-EM tomography observations show that all proto-crowns and all active, double-ring crowns on intact mitochondria have 12-fold (C12) symmetry, implying that the 11-mer (C11) proto-crown rings are artefacts induced by detergent solubilization to release in vivo C12 rings from the outer mitochondrial membrane [14]. Moreover, the atomic- to near atomic-resolution protein A structures derived from these C12 and C11 rings show no significant differences [14]. Reconstruction of the C12 proto-crown structure from negative-stained images revealed a close match to the proto-crown shape defined by cryo-EM tomography and sub-tomogram averaging, into which the ribbon diagram of the C12 protein A multimer structure fit closely (Figure 7A, right panel).

**Figure 7.**
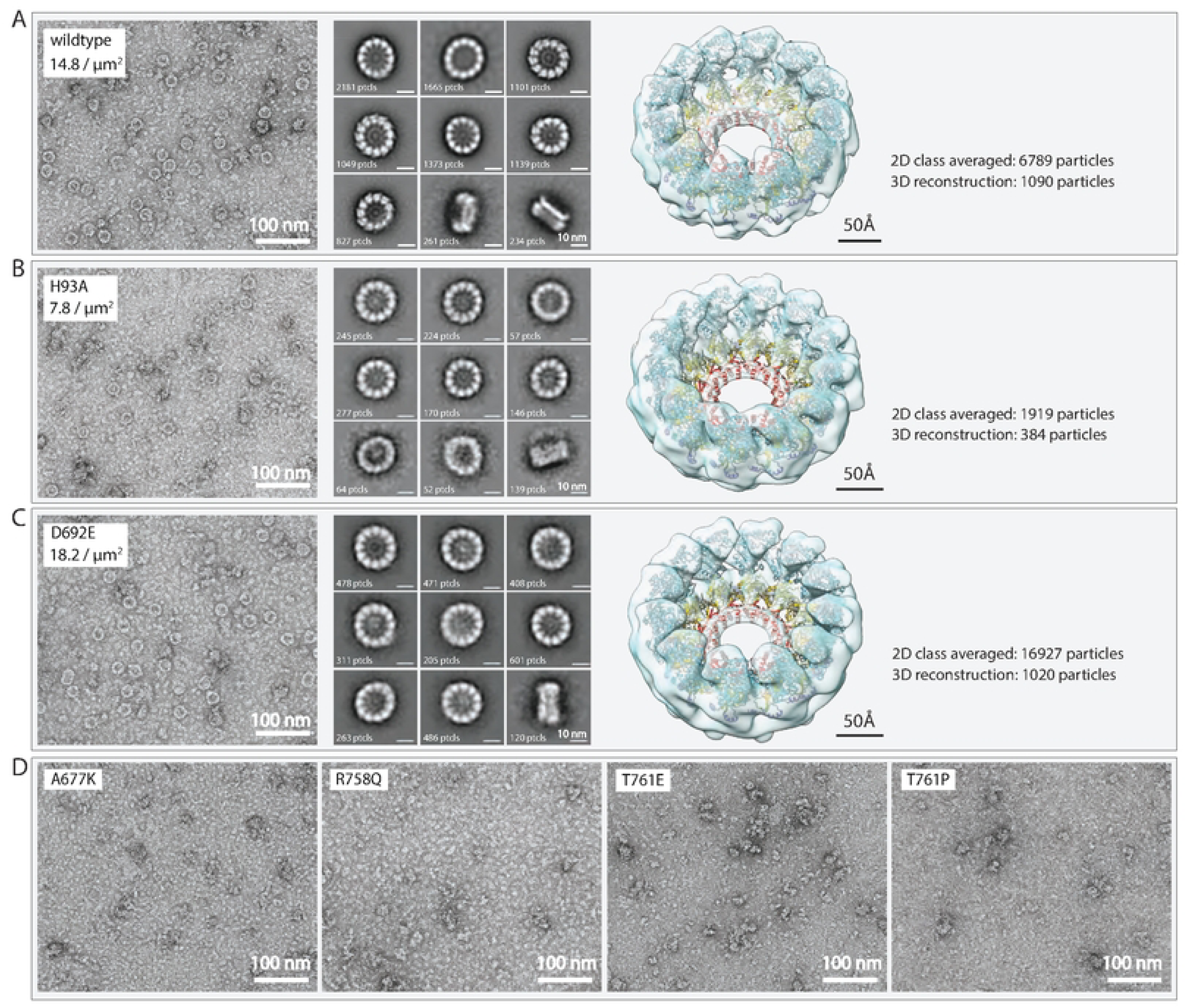
Protein A Pol-Pol interface mutations A677K, R758Q, T761E, and T761P block proto-crown assembly. Negative-stain electron microscopy showed that,while wildtype protein A (A), RNA capping null mutant H93A (B), and RNA polymerase null mutant D692E (C) assemble into lower ring proto-crownsthatcan be purified via baculovirus-launched expression, the A677K, R758Q T761E, and T761P mutants (D) did not assemble into such proto-crowns. 2D Classification (top 9 classes shown) and 3D reconstruction showed that negative-stained H93A and D692E proto-crown samples were essentially indistinguishable from wildtype protein A proto-crowns with any perceived minor differences well below the level of resolution. All reconstructions fit the earlier resolved single particle proto-crown structure (Zhan et al. 2023) equally well.

Parallel assays of extracts from Sf9 cells expressing the protein A H93A capping null mutant or D692E Pol null mutant produced primary grid images, 2D classification analyses and proto-crown reconstructions highly similar to wt protein A, showing that these mutants in catalytic active sites also assembled stable proto-crowns (Figure 7B&C). Any perceived minor differences in the reconstructions were not meaningful as they were below the level of achieved resolution. However, extensive searching of 37, 44, 49, and 39 images from EM grids equivalently prepared from Pol-Pol interface mutants A677K, R758Q, T761E and T761P, respectively, showed no detectable ring structures (Figure 7D). Thus, consistent with their lack of RNA replication, these protein A mutants either failed to form proto-crowns in vivo or their proto-crowns were too unstable to survive isolation. Preparations from cells expressing each of these four mutants did show amorphous, aggregate-like structures in the 10-15 nm size range. Although 2D classification of >300,000 of these particles has not yet revealed any consistent structural subtypes, ongoing studies are exploring if any of these objects might represent intermediate protein A multimers that could further illuminate the structural natures of the assembly defects in these mutants.

### Pol-Pol interface mutations have divergent complementation phenotypes

For more insight into the effects of these proto-crown assembly- and RNA replication-defective Pol-Pol interface mutations, we tested their abilities to complement the Pol and RNA capping defects of D692E and H93A, yielding surprisingly contrasting outcomes (Figure 8). When co-expressed with D692E, A677K gave the highest complementation of any mutant in this report (54-58% of wt protein A RNA replication), T761E gave low level complementation (10-12% of wt protein A) and T761P and R758Q failed to measurably complement. Consistent with their wt RNA capping domains, all four mutants complemented H93A, although to widely differing levels from ∼2 - 30%.

**Figure 8.**
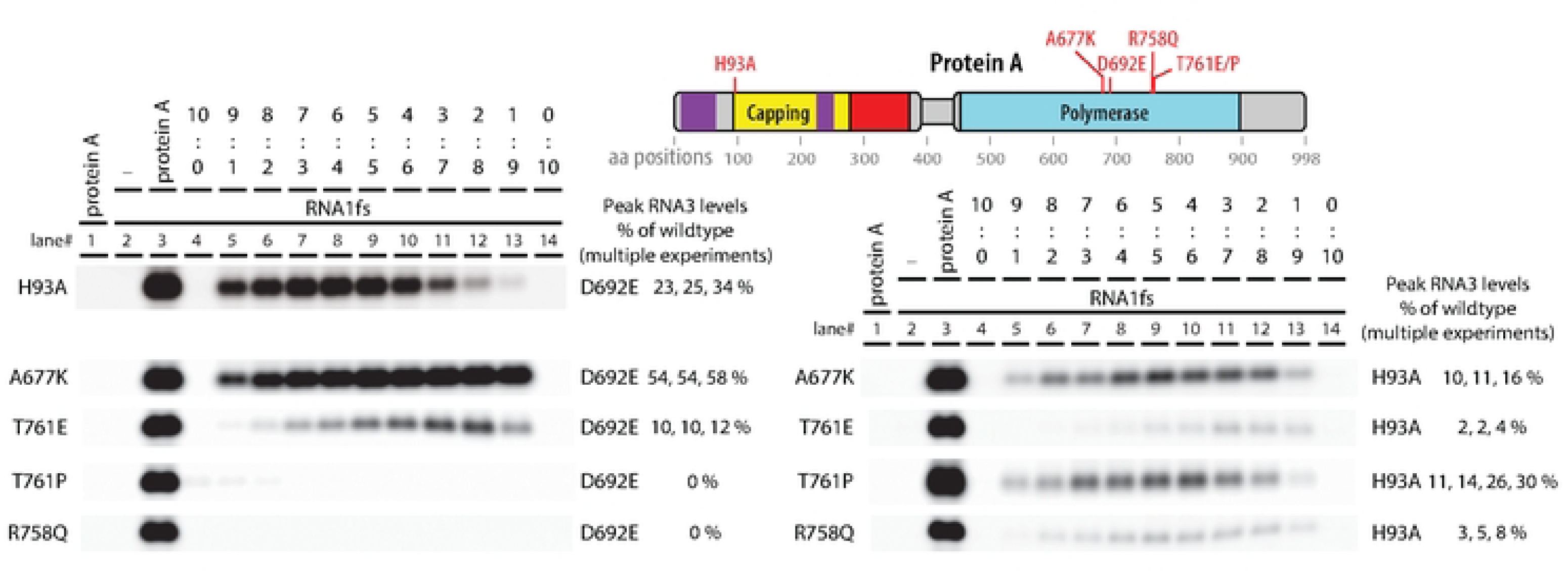
Proto-crown assembly-defective Pol-Pol interface mutants have strikingly divergent phenotypes for complementing RNA capping and Pol null mutants. Complementation assays between the proto-crown assembly mutants and RNA capping H93A or RNA polymerase D692E null mutants were compared to complementation of the H93A with the D692E mutant asareference. Proto-crown assembly mutant A677Kis a fully functional RNA polymerase, but retainsweak RNA capping activity.This is consistent with the idea that the RNA capping functions reside in the proto-crown but that the RNA polymerase functions do not transition through lower proto-crown assembly. Mutants R758Q and T761P abolished such polymerase-providing protein A conformation but likewise retained some RNA capping activity when incorporated in proto-crowns with assist from the H93A mutant to provide wildtype amino acid sequence at the 758 and 761 positions. In contrast to the T761Pmutant, the T761Emutant wasabetter polymerase than it wasacapping enzyme, indicating that the two mutations, even though at the same amino acid position, affected different protein A conformations. RNA3 replication levels were obtained from at least three independent experiments. Representative blots are shown.

Overall, T761E and A677K were similar in each preferentially complementing D692E while complementing H93A to a lower level. In contrast, T761P and R758Q were similar in complementing H93A but not D692E. Thus, even the two substitutions at aa T761 had inverted phenotypes, respectively providing stronger complementation to H93A (T761P) or D692E (T761E). Potential reasons for the failure of T761P and R758Q to complement D692E’s Pol function and for the varying abilities of each mutant to complement H93A are considered in the Discussion.

## DISCUSSION

While recent cryo-EM studies have increasingly refined the structure of nodavirus, CHIKV and coronavirus RNA replication crowns [1,9–12,14,19], understanding of their assembly and operation has remained more limited. This study combined genetic complementation with other approaches to further illuminate the roles and interactions of multi-functional nodaviral protein A in assembly and function of its crown complex and its proto-crown precursor. In addition to other principles, the results reveal in part how nodaviruses have evolved to replicate with high efficiency and wide host compatibility despite an unusually small genome lacking non-structural protein processing, through divergent uses of full-length, multi-domain protein A in multiple steps, functions, conformations and crown sites.

### Complementation shows functional separability of nodaviral RNA polymerase and capping

Active site protein A mutants H93A and D692E respectively block RNA capping and RNA polymerase activity, but each assemble stable proto-crowns (Figure 7) and together efficiently co-immunoprecipitate (Figure 2B). Importantly, when co-expressed, capping null mutant H93A and Pol null mutant D692E complemented each other’s defects to support RNA replication over a wide range of plasmid ratios (Figures 2B, 3A, 8). Similarly, D141A, which blocks a distinct capping step, also strongly complemented D692E (Figure 3B).

Thus, RNA polymerase and RNA capping ability are not required in all copies of protein A, nor together in any single copy of protein A. Rather, these activities are functionally separable and can – and perhaps must – be provided by different copies of protein A. This separable, cooperative action is consistent with the presence in nodavirus crowns of multiple protein A conformations (Figure 1DE), and with a priori considerations and structural conservation between nodavirus and alphavirus crowns noted in the Introduction, implying that 5’ capping of nascent (+)RNAs is provided by capping domains in the crown floor, while Pol activity for that (+)RNA synthesis is provided by a polymerase domain at the center of that floor (Figure 1E).

Such division of enzymatic function between alternate protein A positions and conformations in the crown should place limits on the efficiency of complementation between H93A and D692E. For example, if co-expression were equal and assembly unbiased, half of crowns would have the central Pol position occupied by H93A (Pol+) and be functional for Pol activity, while the other half would bear D692E (Pol-) and be nonfunctional, limiting complementation to ∼50% of RNA replication levels supported by wt protein A. Similar issues apply to RNA 5’ capping. Currently it is not known how the active Pol transfers a newly synthesized 5’ (+)RNA end to an RNA capping domain (Figure 1DE). Nonetheless, in complementation between H93A and D692E, RNA accumulation may be further reduced by non-productive capping interactions due to mixed occupancy of the crown floor by capping-inactive H93A and capping-active D692E protein A variants. Interestingly, these considerations of mixed Pol and capping occupancy suggest that the observed complementation of H93A or D141A with D692E, which reached 25-40% of wt protein A RNA replication (Figure 3AB, 8), might be close to theoretical maximum levels.

### Key RNA capping steps are not separable between two copies of protein A

Nodavirus and alphavirus RNA capping domains share a common protein fold, unusual capping pathway and formation of 12-mer rings [1,11,14,27]. Assembly of these 12-mer rings is essential to activate RNA capping [26,27]; our unpublished results], raising a possibility of cooperative action. However, D141A and H93A capping mutants, respectively incapable of GTP N7 methylation and m^7^GMP esterification, did not complement each other (Figure 3C). Thus, m^7^GTP synthesized by H93A was unable to diffuse to adjacent D141A subunits for covalent linkage to aa H93, experimentally confirming that – although the capping domain only functions as a 12-mer - these successive steps must occur in a single capping subunit.

### RNA polymerase and capping require full protein A contexts

Successful complementation between protein A derivatives bearing distinct inactivated domains (Figures 2 and 3) prompted testing if such complementation might tolerate inactivation by selected deletions as well as point substitutions. However, initial analysis with 100 aa overlapping deletions tiled across protein A showed that essentially all deletions abolished complementation with either H93A or D692E (Figure 4B). More targeted deletions further showed that, despite marked differences between membrane contacts of the apical and basal protein A rings (Figure 1D), both of protein A’s distally encoded membrane interaction domains were also required to complement H93A or D692E (Figure 5).

While it seems likely that some complementation pairings would tolerate specific smaller deletions – e.g., structure-guided deletions of particular peripheral loop regions – the strongly negative results from partial deletions across the entire length of protein A imply that Pol, capping and likely other protein A functions have evolved to function in the context of full-length protein A. This is consistent with our prior findings that >95% of protein A is full length and that the bulk of the crown is built from 24 full-length protein A’s that in alternate conformations form the apical and basal rings (Figure 1DE) [14]. In these contexts, many deletions even in inactivated domains could interfere with crown assembly and structure.

The potential impact of such deletions was less obvious for the central density strongly implicated as the active Pol domain, since from current structural data it is not yet clear how or even if this central putative Pol is connected to flanking protein A sequence (see Introduction). If, e.g., this central density was a free polymerase domain produced by proteolysis or alternate expression, parts of the capping and/or central channel domains might well have been dispensable for complementation with D692E. That none were dispensable favors alternate hypotheses linking this central density more directly to full length protein A, such as a Pol domain “bungee jumping” from the apical or basal rings on its long flexible linker to the N-proximal capping/central channel region (Figure 1A; see also Introduction).

The single deletion that showed any complementing activity was Δ897-998, which weakly complemented H93A (Figure 4B). Amino acids 897-998 comprise a C-terminal intrinsically disordered region. Unlike the more balanced complementation between H93A and D692E (Figure 3A), in this case optimum complementation favored excess H93A (Figure 4C), showing that RNA replication was more sensitive to losing copies of this C-terminal domain than to losing capping or polymerase activity. Intrinsically disordered regions of proteins are increasingly recognized as key interaction regions that in this case might be required for condensation of viral replication proteins by liquid-liquid phase separation, crown assembly interactions, host factor recruitment, or other functions [31–33].

### Pol-Pol interface mutants are defective in proto-crown assembly and vary in complementation abilities

The proto-crown, the earliest known crown assembly intermediate, is primarily stabilized by extensive contacts between its floor subunits, while its ringed Pol subunits (Figure 1E) interact much more weakly [14]. Nevertheless, when we targeted mutations into loops at the Pol-Pol interface and identified four that blocked RNA replication (Figure 6), all four markedly inhibited assembly of stable proto-crowns (Figure 7). Thus, in addition to the core foundation of extensive floor interactions, compatible junctions between Pol domains are also necessary for proto-crown stability and are potential targets for viral interference.

Complementation assays revealed further impacts of these Pol-Pol interface mutations that were highly varied (Figure 8). Proto-crown assembly impaired A677K complemented D692E unusually well, showing that A677K’s Pol domain must be highly active and readily assemble into the presumably central position required for its function, without transitioning through a lower proto-crown ring assembly. The reduced ability of capping-impaired A677K to assemble into the crown floor sites linked to capping will also leave these positions to be predominantly occupied by capping-active D692E, further enhancing complementation efficiency. In contrast, T761P and R758Q failed to complement D692E at all. Proline substitution is often disruptive to protein folding, with larger scale structural effects that in T761P may well block Pol enzyme function and/or assembly. Moreover, unlike substitution-tolerant positions A677 and T761, all tested substitutions at R758 blocked replication and all but R758Q blocked all complementation (Figure 6). Thus, position R758 is highly restricted, perhaps to satisfy separate constraints for more than one protein A function or conformation.

Conversely, all four Pol interface mutants complemented H93A to some degree, reflecting that all retain wt RNA capping domains. Their varied levels of complementing H93A seem likely due to varied abilities to assemble into the crown floor where capping occurs – due to variations in intrinsic defects in proto-crown assembly (Figure 7) or possibly to partial complementation of some mutants’ assembly defects by the wt Pol interface of H93A. These diverse results reflect multi-functional protein A’s interrelated enzymatic, assembly and other abilities and its conformational isomerism, and illustrate how complementation synergizes with other results to uncover the underlying mechanisms.

## METHODS

### Recombinant DNA plasmid construction

Standard molecular cloning procedures were used to place FHV RNA1 or protein A-encoding and derivative mutant sequences under control of the baculovirus IE1 promoter for expression in *Drosophila* S2 cells or the baculovirus polyhedrin promoter in pOET1 transfer plasmids needed to generate recombinant baculoviruses. Plasmid pIE1hr5-FHVRNA1fsRz-D692E was used to launch an FHV RNA1 replication template containing an early protein A ORF frameshift and, as an additional precaution, a catalytic site inactivating mutation in the RNA-dependent RNA polymerase domain. Precise 3’ end termination was mediated by a downstream hepatitis delta virus ribozyme sequence. Plasmid pIE1hr5-FHVptnA(rr) and derivatives were used to express FHV protein A and mutants. These protein A expression plasmids contained the protein A ORF but did not have the authentic FHV RNA1 5’ and 3’ ends. The region previously shown to be important for recruitment of RNA1 into replication complexes [17] was recoded with translationally silent mutations to preserve the correct FHV protein A amino acid sequence. Protein A mutant plasmids were generated by PCR using mutation-introducing oligonucleotides, or by replacement of restriction enzyme fragments with synthetic mutant DNA gBLOCKs purchased from either IDT DNA Technologies (Coralville, IA, USA) or TWIST Bioscience (San Francisco, CA, USA). Plasmid sequences were verified using Sanger sequencing at the UW-Madison Biotechnology Center Core Sequencing Facilities or at Functional Biosciences, Madison WI.

### Recombinant baculoviruses

Recombinant baculoviruses expressing FHV protein A or its mutant derivatives were generated using pOET1-based protein A-expressing transfer plasmids with the FlashBac system (Mirus-Bio, Madison, WI, USA) according to provided instructions. Baculovirus titers were determined by Express Systems, Inc (Lake Forest, CA, USA).

### Cell culture and DNA transfection

*Drosophila melanogaster* S2 cells were maintained at 28°C in either Schneider’s medium supplemented with 120U/ml penicillin, 100 µg/ml streptomycin and 2mM L-glutamine (P/S/G, GIBCO/ThermoFisher, Waltham, MA, USA) and 10% inactivated bovine calf serum ( BCS, Cytiva/HyClone, Logan, UT, USA) or in ExpressFive medium (GIBCO/ThermoFIsher) supplemented with P/S/G. *Spodoptera frugiperda* Sf9 cells were grown under the same conditions in SF900 medium (GIBCO/ThermoFisher) containing P/S/G. For plasmid transfection, 2.5 x 10^6^ Drosophila S2 cells/well were seeded in 6-well plate format. The next day, prior to transfection, culture medium was replaced with 1 ml Schneider’s medium supplemented with P/S/G and 10% BCS. DNA plasmid transfection mixtures in 100 µl unsupplemented Opti-MEM (GIBCO/ThermoFisher) containing 1 µg RNA1 template plasmid and 500 ng protein A expression plasmid (single or mixed) were complexed with 6 µl *Trans*IT-insect reagent (Mirus Bio, Madison, WI, US) for 15-20 minutes at room temperature and added to the cells. Transfections were incubated at 28°C, and after 1 hour provided with another 500 µl of Schneider’s medium supplemented with P/S/G and 10% BCS. Cells were harvested for protein and RNA at 65-70 hours post-transfection.

### RNA analysis

Total RNA was extracted from half of the transfected cells using a Maxwell RSC48 automated RNA extraction system and reagents (Promega, Madison, WI, USA). For RNA analysis, northern blots were used with biotinylated RNA probes essentially as described previously [34]. RNA probes detecting part of the RNA1 and RNA 3 common sequence were generated using nucleotide concentrations at 0.5 mM for ATP, CTP, and GTP, 0.3 mM for UTP/0.2 mM Bio-16 UTP. Blots with hybridized biotinylated probes were visualized using IRDye680RD-streptavidin or IRDye800CWstreptavidin and imaged using a LI-COR Odyssey system RNA levels were quantified using LI-COR Image Studio software (version 5.2).

### Protein Analysis

Total protein was extracted from half of the transfected cultures for western blotting using primary antibodies against FHV protein A and tubulin (Sigma, T9026) each at 1:2,000 dilution as described previously [34]. Westerns confirmed protein A expression for all experiments presented. Co-immunoprecipitation experiments were carried out essentially as described previously [34], but using anti-FLAG (M2) antibody-conjugated beads (Sigma, A2220-1ml) for the precipitation and chicken anti-MYC antibody (Bethyl Laboratories, A190-103A at 1:1,000 dilution) for the western.

### Isolation of mitochondria

In 20 ml SF900 medium supplemented with P/S/G, 5 x 10^8^ Sf9 cells were infected with recombinant baculovirus expressing alfa-tagged FV protein A or mutant derivatives at an m.o.i. of 5-10. After 1 h of shaking at 120 rpm at 28°C, culture volumes were increased to 400 ml and further incubated for 3 days. Small aliquots of the cultures were set aside for protein expression analysis on western blots, while the bulk of the cultures were used for mitochondrial isolation using Qiagen’s Qproteome kit. Cells were pelleted for 5 min at 500 x g, resuspended in 20 ml PBS, pelleted, resuspended in 8 ml 0.9% (w/v) NaCl, pelleted again, and resuspended in 2.5 ml Qproteome lysis buffer and 100 µl 25X Roche protease inhibitor. Samples were rocked for 10 min on ice and centrifuged at 1000 x g for 10 min at 4°C. Supernatants containing cytosol were discarded and pellets were resuspended in 2.0 ml Qproteome Disruption buffer with 80 µl 5x Roche protease inhibitor. Lysates were slowly drawn up and down 10 times through a 25 gauge syringe, taking care to avoid bubbling. Lysates were centrifuged at 1000 x g for 10 min at 4°C and turbid-appearing supernatants were transferred to 2 ml centrifuge tubes, centrifuged at 6000 x g for 10 min at 4°C. Visible pellets containing mitochondria were resuspended in 750 µl Qiagen Mitochondria Purification Buffer and carefully layered on top of another 750 µl Qiagen Mitochondria Purification Buffer, underlaid earlier with 500 µl Disruption Buffer. Samples were centrifuged at 14,000 x g for 15 min at 4°C. 1.5 ml of supernatants were carefully removed to not disturb the mitochondrial pellet which were then resuspended in the remaining solution and transferred to a new 2 ml tube of pre-determined weight. Using multiple rounds of repeated centrifugation, 1.5 ml Mitochondria Storage Buffer was added, samples were centrifuged at 8000 x g for 10 min at 4°C, and 1.5 ml of supernatant was carefully removed, until mitochondria had formed a pellet at the bottom of the tube.

### Crosslinking and solubilization of protein A / proto-crowns from mitochondria

Mitochondrial pellets were resuspended to a final concentration of 45 mg/ml in 1x RSB Hypo Buffer (a 2:3 mixture of 2.5x MS Homogenization Buffer and 1x RSB Hypo Buffer (10 mM HEPES pH 7.5, 10 mM NaCl, 0.2 mM EDTA) containing 2.5 mM Disuccinimidyl glutarate (DSG). Samples were gently rotated for 30 min at 4°C. Cross-linking was quenched by adding 1M Tris-HCl pH 8 to a final concentration of 25 mM and samples were further rotated for 15 min at room temperature. An equal volume of 4% n-Dodecyl-B-D-maltoside (DDM) Solubilization Buffer (20 mM Tris, 300 mM NaCl, 2 mM MgCl_2_, 4% DDM, 4% glycerol, 1x Roche protease inhibitor, 25U/ml benzonase) was added to the cross-linked mitochondria. Samples were gently rotated for 30 min at room temperature and centrifuged at 21,000 x g for 15 min at 20°C. Supernatants with solubilized alfa-tagged protein A / proto-crowns were collected.

### ALFA-tag mediated protein A / proto-crown purification and concentration

100 µl (50 µl bed volume) ALFA Selector PE 50% slurry resin (NanoTag Biotechnologies, Gottingen, Germany) spin) was equilibrated in 200 µl Wash Buffer (0.05% DDM, 20 mM Tris pH 8.0, 150 mM NaCl, 1% glycerol) and pelleted at 21,000 x g. Wash buffer was removed, solubilized proto-crowns were added and the mixture gently rotated for 30 min at room temperature. Samples were loaded onto a Bio-Rad Poly-Prep column (2 ml bed volume and 10 ml reservoir) allowing the resin to pack. After release of flow-through, the ALFA-PE resin was washed with 600 µl (12 bed volumes) Wash Buffer. ALFA-tagged protein A / proto-crowns were eluted from the PE resin in 100 µl (2 bed volumes) Elution Buffer (Wash buffer containing 1µl 100 x ALFA-peptide) by nutating for 25 min at room temperature during which time samples were mixed every 5 min. Eluates were collected in “low-binding” 1.5 ml tubes and concentrated to ∼30 µl final volume using Vivaspin500 50 kDa cut-off PES columns (pre-rinsed with 400 µl Wash buffer) by sequential centrifugation for 2 min at 4°C using 400 x g increments starting at 2400 x g, typically ending at 4,400 x g. Concentrates were either immediately used for negative staining on EM grids or stored in 1.5 ml Low-Bind tubes at −80°C.

### Negative staining, EM-imaging, particle classification and 3D reconstruction

3 µl of concentrated proto-crown preparations were spotted onto copper grids (EMS #CF300-Cu-50) that within an hour prior to use had been glow-discharged using a PELCO easiGLow for 30s at 20 mA (Ted Pella, Inc., Redding, CA , USA). After 1 minute, grids were side-blotted a on Whatman filter paper (Cytava #1002-090) and washed and blotted twice and stained and blotted twice in 20 µl drops of wash solution (0.05% DDM, 20 mM Tris pH 8.0, 150 mM NaCl, 1% glycerol), and 2% uranyl acetate (UA, EMS #22400-2), respectively, and finally left floating face-down on a drop of 2% UA for 1 minute before slowly blotting on top of the previously discarded UA. Grids were air-dried for 1 minute and stored in grid boxes in a vacuum desiccator until ready for EM imaging at 45,000x magnification with pixel size of 2.646Å using a Talos F200C Transmission Electron Microscope equipped with a OneView Camera (ThermoFisher). Images were collected using SerialEM software [35]. Electron micrographs were imported into CryoSPARC v4.7.1 [36] for processing, following the parameters described in the CryoSPARC tutorial (https://guide.cryosparc.com/processing-data/tutorials-and-case-studies/negative-stain-data) and the standard negative staining protocol [37]. After CTF estimation [38], manual curation was performed to remove poor-quality images. The blob picker was used to identify particles with a maximum diameter of 20 nm and a minimum diameter of 10 nm, and the picking results were inspected. Several rounds of 2D classification were carried out to eliminate poor-quality particles, followed by *ab initio* reconstruction. Subsequent homogeneous refinement with C12 symmetry applied produced the final 3D average maps. The number of particles used for 2D classification and 3D reconstruction are provided with the representation of the results in Figure 7.

## ACKNOWLEDGMENTS

We thank other members of our group at the John and Jeanne Rowe Center for Research in Virology and colleagues in the larger Morgridge Institute for Research and UW-Madison community for valuable discussions. We also thank Timothy Grant, Amy Strydom, the UW-Madison 3D EM Core Facility and Cryo-EM Research Center (CEMRC) for support in electron microscopy and data collection, analysis and interpretation, and Sohini Sengupta for laboratory assistance.

Part of our analyses were performed using the resources and assistance of the UW-Madison Center for High Throughput Computing (CHTC) in the Department of Computer Sciences. The CHTC is supported by UW-Madison, the Advanced Computing Initiative, the Wisconsin Alumni Research Foundation, the Wisconsin Institutes for Discovery, and the NSF, and is an active member of the Open Science Grid which is supported by the NSF and the US Department of Energy’s Office of Science. P.A. gratefully acknowledges support from the John and Jeanne Rowe Center for Research in Virology at the Morgridge Institute for Research and the University of Wisconsin - Madison Steenbock Professorship in Microbiology.

